# Sex differences in binge alcohol drinking and the behavioral consequences of protracted abstinence in C57BL/6J mice

**DOI:** 10.1101/2023.05.12.540565

**Authors:** Jean K. Rivera-Irizarry, Lia J. Zallar, Olivia B. Levine, Mary Jane Skelly, Jared E. Boyce, Thaddeus Barney, Ruth Kopyto, Kristen E. Pleil

**Affiliations:** Neuroscience Graduate Program, Weill Cornell Graduate School of Medical Sciences, Weill Cornell Medicine, Cornell University, New York, NY, USA; Pharmacology Graduate Program, Weill Cornell Graduate School of Medical Sciences, Weill Cornell Medicine, Cornell University, New York, NY, USA; Department of Pharmacology, Weill Cornell Medicine, Cornell University, New York, NY, USA

## Abstract

**Background:** Binge alcohol drinking is a risk factor linked to numerous disease states including alcohol use disorder (AUD). While men binge drink more alcohol than women, this demographic gap is quickly shrinking, and preclinical studies demonstrate that females consistently consume more alcohol than males. Further, women are at increased risk for the co-expression of AUD with neuropsychiatric diseases such as anxiety and mood disorders. However, little is understood about chronic voluntary alcohol drinking and its long-term effects on behavior. Here, we sought to characterize sex differences in chronic binge drinking and the effects of protracted alcohol abstinence on anxiety- and affective-related behaviors in males and females.

**Methods:** We assessed binge alcohol drinking patterns in male and female C57BL/6J mice using a modified Drinking in the Dark (DID) paradigm in which mice received home cage access to one bottle of 10% or 20% alcohol (EtOH) or water for 2 hrs per day on Days 1-3 and to two bottles (EtOH/H2O + H2O) for 24 hrs on Day 4 for eight weekly cycles. Mice were then tested for the effects of protracted abstinence on avoidance, affective, and compulsive behaviors.

**Results:** Female mice consumed more alcohol than males consistently across cycles of DID and at 2, 4, and 24-hr timepoints within the day, with a more robust sex difference for 20% than 10% EtOH. Females also consumed more water than males, an effect that emerged at the later time points; this water consumption bias diminished when alcohol was available. Further, while increased alcohol consumption was correlated with decreased water consumption in males, there was no relationship between these two measures in females. Alcohol preference was higher in 10% vs. 20% EtOH for both sexes. During protracted abstinence following chronic binge drinking, mice displayed decreased avoidance behavior (elevated plus maze, open field, novelty suppressed feeding) and increased compulsive behavior (marble burying) that was especially robust in females. There was no effect of alcohol history on stress coping and negative affective behaviors (sucrose preference, forced swim test, tail suspension) in either sex.

**Conclusion:** Female mice engaged in higher volume binge drinking than their male counterparts. Although females also consumed more water than males, their higher alcohol consumption was not driven by increased total fluid intake. Further, the effects of protracted abstinence following chronic binge drinking was driven by behavioral disinhibition that was more pronounced in females. Given the reciprocal relationship between risk-taking and alcohol use in neuropsychiatric disease states, these results have implications for sex-dependent alcohol drinking patterns and their long-term negative neuropsychiatric/physiological health outcomes in humans.

**Summary:** The overconsumption of alcohol is a widespread public health issue linked to numerous diseases and mental health issues including anxiety. Binge drinking, defined as having 4-5 drinks in a 2-hour period, is more common in men than women but that demographic gap is shrinking. Mice are commonly used as an animal model of alcohol consumption and binge drinking to study behaviors associated with/resulting from alcohol consumption. We found that female mice consumed more alcohol compared to their male counterparts. While female mice also drank more water than males, high alcohol consumption was not correlated with water consumption in females. In addition, following long-term alcohol consumption and protracted abstinence, mice displayed behavioral disinhibition marked by reduced avoidance and increased compulsive behavior; this phenotype was pronounced in females. As reduced adaptive anxiety/increased risk-taking behavior and alcohol consumption can promote one another, our results suggest that women may be especially vulnerable to the negative outcomes associated with chronic alcohol drinking.

**Highlights:**

- Female mice consistently consumed more alcohol per bodyweight than males, with a more robust sex difference for high (20%) than low (10%) concentration alcohol.
- Females consumed more water than males at later (4-hr and 24-hr) but not early (2-hr) timepoints, and this effect was diminished when alcohol was available.
- Higher alcohol consumption was correlated with decreased water consumption in males but not females, suggesting that females’ greater alcohol consumption is not due to higher total fluid intake.
- Alcohol preference was higher for 10% versus 20% alcohol in both sexes.
- Mice in protracted abstinence (2-6 wks) from binge alcohol drinking displayed a reduction in avoidance behavior and increase in compulsivity compared to water-drinking controls, especially in females. There was no effect of protracted alcohol abstinence on anhedonia in either sex.

## Introduction

Excessive alcohol use is a critical public health issue with wide ranging social and health-related negative consequences. Alcohol use disorder (AUD), which affects nearly 30 million people in the United States alone (1), is highly co-expressed with other neuropsychiatric diseases including anxiety and mood disorders (2, 3). The negative outcomes associated with alcohol are primarily driven by the pattern and cumulative volume of consumption (3). In particular, binge alcohol drinking, defined as the consumption of alcohol resulting in a blood ethanol concentration (BEC) of >0.08 mg/dl, is one of the most significant predictors of later development of AUD and other co-morbid neuropsychiatric diseases (2, 4). Indeed, persistent cycles of binge drinking to intoxication followed by withdrawal persists among individuals with AUD (3, 5). Binge drinking and AUD are more common among men than women, although this gap in prevalence has been shrinking in recent years as women’s alcohol drinking is increasing at a faster rate than men’s (3, 6). Current epidemiological data show that adolescent girls are already as likely to express AUD as adolescent boys (7). Females in other mammalian species including rodents drink as much or more than their male counterparts when intake is adjusted to bodyweight (8-12). Nonetheless, the complexities of binge drinking behavior remain poorly understood in females, including in rodent models such as the commonly used C57Bl/6J mouse strain that has been employed robustly in males. In particular, it is unclear whether females consume more than males due to differences in alcohol preference or motivated voluntary consumption, as the studies performed to date report a mix of results showing elevated female preference for at least some time points of alcohol exposure (10, 11) or no sex differences (9, 13-15), depending on the alcohol exposure paradigm and length.

Alcohol drinking behavior significantly contributes to the expression of other maladaptive behaviors associated with neuropsychiatric diseases, such as anxiety, anhedonia, and compulsive behaviors. Studies have found that chronic alcohol exposure has divergent effects on behavior depending on rodent species/strain, alcohol administration paradigm, and state of alcohol dependence or duration of abstinence (recently reviewed in detail by Bloch and colleagues (16)). The effects of protracted abstinence (<1 week) from chronic (4 weeks+) alcohol exposure on emotion-related behaviors in male C57BL/6J mice varies widely, with some showing increased avoidance, anhedonia, and compulsive behaviors (17-22) and others reporting no effects in some or all measures (10, 20, 23). However, these studies vary widely in the alcohol exposure paradigms, durations of alcohol exposure, and behavioral testing time points employed. Relatively little is known about the behavioral effects of chronic voluntary binge alcohol drinking beyond a week into abstinence. Furthermore, while women are at an increased risk of co-expression of AUD with anxiety and affective disorders (24-26), there is a dearth of females used in the pre-clinical literature exploring the effects of protracted alcohol exposure on behavior, using many common protocols including vapor inhalation and voluntary binge drinking models such as intermittent access (IA) and Drinking in the Dark (DID; 16). As such, the field has not adequately assessed the role of sex in the relationship between alcohol and emotional states.

Here, we investigated sex differences in binge alcohol consumption, pattern, and preference using a modified DID paradigm, as well as the effects of chronic binge alcohol drinking and protracted abstinence on stable behavioral phenotypes. We found that females consumed more alcohol than their male counterparts at all timepoints (2, 4, and 24 hrs) and more water than males at later timepoints (4 and 24 hrs) only when alcohol was not available. Intriguingly, higher alcohol consumption was correlated with reduced water consumption in males but not females, suggesting that females’ greater alcohol consumption than males was not due to a generally higher intake of fluid. Additionally, we found that compared to water-drinking controls, alcohol-exposed mice displayed a consistent reduction in avoidance behavior during protracted abstinence (2-4 weeks post-alcohol) and increased compulsive-like behavior without alterations in affective behaviors. Moreover, these aberrant risk-taking behaviors were especially pronounced in females. As females consumed higher levels of alcohol across the eight weeks of exposure, these results may reflect increased susceptibility to aberrant plasticity in females and/or a dose-dependent effect of alcohol.

## Methods

### Animals

Male and female C57BL/6J mice were purchased from Jackson Laboratories (stock #00064) at eight weeks of age. Mice were housed in same-sex cages of five on a reverse 12:12 hr light cycle colony room with lights off at 7:30 am daily, given *ad libitum* access to water and standard chow, and allowed to acclimate for at least one week. Mice were singly housed one week prior to the beginning of binge alcohol (20% or 10% EtOH) or water (H2O) drinking. Following eight weekly cycles, 20% EtOH and H2O groups underwent a battery of behavioral tests over six weeks of abstinence. All behavioral testing occurred during the dark phase of the light-dark cycle, beginning approximately three hrs into the start of the dark phase. For all tests occurring outside of the home cage, mice were habituated to the testing room for one hr before behavioral testing, and assays were conducted under 250 lux lighting conditions and analyzed using Ethovision XT (Noldus). All experimental procedures were approved by the Institutional Animal Care and Use Committees at Weill Cornell Medicine.

### Binge alcohol drinking

We used a modified version of the standard DID (27) procedure, allowing us to assess alcohol preference within the DID paradigm across eight weekly cycles (**Fig. 1a**). Home cage water bottles were replaced with 50 ml bottles containing alcohol (EtOH, 20% or 10%) or water (H2O) as a control three hrs into the dark phase of the light cycle for four consecutive days each weekly DID cycle. On Mon-Wed, mice received two hours of access, as in the standard DID paradigm (8). On Thurs, mice received a modified access to two bottles (one EtOH and one water for EtOH drinking groups, and two water bottles for H2O DID) for 24 hrs, and bottle weights were taken 2 hr, 4 hr, and 24 hr into access. EtOH/water bottles were then replaced with home cage water bottles on Fri morning until the following DID cycle, which began 72 hr later the following Mon. In the latter half of the paradigm, a subset of 10% EtOH mice underwent unsuccessful fear conditioning (due to technical issues that prevented the floor grid from providing shocks) before cycles 5 and 8; because these mice did not differ from their counterparts in any measures recorded, all 10% EtOH DID mice are included within as a comparison to 20% EtOH and H2O DID mice for alcohol drinking. Throughout, *intake* is defined as raw EtOH/H2O/fluid while *consumption* is defined as EtOH/H2O/fluid intake normalized to bodyweight. Following eight cycles, the 20% EtOH and H2O DID cohorts underwent a battery of behavioral tests (**Fig. 6a**).

**Figure 1:**
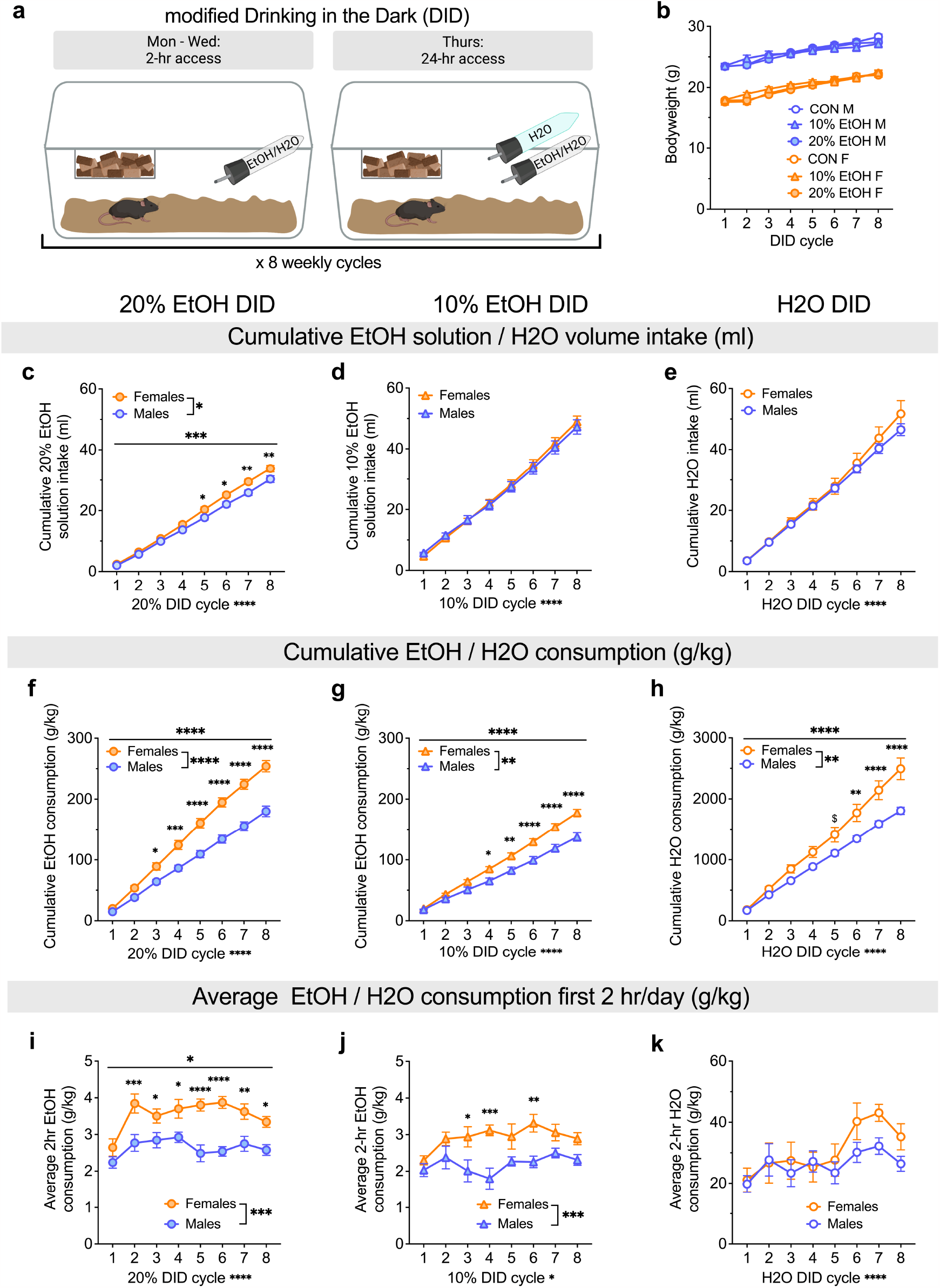
Alcohol and water consumption are sex- and alcohol concentration-dependent. **A)** Schematic depicting the modified Drinking in the Dark (DID) binge drinking paradigm in C57BL/6J mice. **B)** Bodyweight gain was similar across DID cycles in H2O, 10% EtOH, and 20% EtOH DID mice of both sexes. **C)** Cumulative intake of 20% EtOH solution (ml) was higher in females compared to males on DID cycles 5-8. **D)** There was no sex difference in cumulative 10% EtOH intake (ml) across DID cycles. **E)** There was no sex difference in cumulative H2O intake (ml) across DID cycles. **F)** Cumulative EtOH consumption normalized to bodyweight (g/kg) was higher in females compared to males on DID cycles 3-8. **G)** Females consumed more 10% EtOH (g/kg) compared to males on DID cycles 4-8. **H)** Females consumed more H2O (ml/kg) compared to males during DID cycles 6-8, with a trend on cycle 5. **I)** Females displayed higher average 20% EtOH consumption (g/kg) during the first two hrs/day of access on DID cycles 2-8. **J)** Females displayed higher average 10% EtOH consumption (g/kg) on the first two hrs/day of access on DID cycles 3, 4, and 6. **K)** There was no difference between males and females on average H2O consumption during the first two hrs/day of water access. *P < 0.05, **P < 0.01, ***P < 0.001, ****P < 0.0001 in 2xRM-ANOVA main effects and interactions of sex and DID cycle, as well as post hoc t-tests with H-S corrections between M and F. ^$^P < 0.10 for post hoc t-tests with H-S corrections between M and F.

### Sucrose preference test

Seventy-two hr after the last alcohol/water access of eight-cycle DID (20% EtOH and H2O), mice were assessed for their preference for 1% sucrose over three consecutive days, followed by 2% sucrose for the subsequent three days, using a 24 hr two-bottle choice assay as previously described (28). The home cage water bottle was replaced with two 50 ml bottles, one containing 1% sucrose and one containing water, and bottles were weighed and replaced in switched positions every 24 hr for three days. The following three days, the same procedure was repeated with bottles containing water and 2% sucrose. Sucrose and water intake were recorded, and consumption and preference were analyzed for days 2-3 for each sucrose concentration. Data were excluded if a bubble in the bottle spout prevented access to the sucrose on that day.

### Open field test

Approximately two weeks following the last alcohol exposure, mice were tested for avoidance behavior and locomotion in the open field test (OF) as previously described (28). The test was conducted in a plexiglass arena (50 × 50 × 34.5 cm) with a gray floor and each mouse was placed in one corner of the arena and allowed to freely explore for 30 min. Total time spent in the center of the arena (all four paws in the 25 cm × 25 cm area in the center of the arena) and periphery were quantified to calculate the percent center time. The total distance traveled in the maze (m) was used to measure locomotion while the percent time in the center of the maze was used to assess avoidance-like behavior.

### Elevated plus maze

Approximately 2.5 weeks following the last alcohol exposure, the elevated plus maze (EPM) test was conducted as previously described (28) in a plexiglass maze with two open and two closed arms (35 cm l × 5.5 cm w, with 15 cm h walls for closed arms) extending from a central platform (5.5 cm ×5.5 cm) elevated 50 cm above the floor. Mice were placed in the center of the EPM facing an open arm and allowed to freely explore the maze for five min. The percent time spent in the open arms was used to assess avoidance behavior and ambulatory locomotion was also analyzed. Due to a technical issue, EPM data for some mice could not be analyzed and were therefore excluded.

### Light-dark box

Approximately three weeks following the last alcohol exposure, the light-dark box (LDB) test was conducted as previously described (28) in a rectangular box divided into two equal compartments (20 cm l × 40 cm w × 34.5 cm h): one dark with a closed lid and the other with an open top and illuminated by two 60 W bulbs placed 30 cm above the box. The two compartments were separated by a divider with a 6 cm x 6 cm cut-out passageway at floor level. At the beginning of a trial, each mouse was placed in a corner of the light compartment and allowed to freely explore the apparatus for 10 min. The number of light side entries and total time spent in the light compartment as compared to the dark compartment were used to assess avoidance behavior.

### Marble burying task

Approximately 3.5 weeks following the last alcohol exposure, mice were placed in a clean, standard rat cage (26 cm l x 48 cm w x 20 cm h) with 5 cm of fresh, unscented bedding containing 20 evenly-spaced marbles. Following 30 min, the number of marbles buried, with a threshold of at least ⅔ of the marble being covered by bedding, was assessed by the experimenter and recorded.

### Novelty suppressed feeding

Approximately four weeks following the last alcohol exposure, mice underwent the novelty suppressed feeding (NSF) test. Mice were exposed to a fruit loop in their home cages for two consecutive days, then fasted overnight for 18-20 hrs prior to testing. During the assay, the mouse was placed into a novel environment (plexiglass arena; 50 × 50 × 34.5 cm) containing fresh bedding and a fruit loop on a piece of weighing paper in the center of the arena. The latency to retrieve and bite the fruit loop was recorded by an experimenter. As soon as the mouse began to bite the fruit loop, or 5 min had elapsed, the mouse was removed and placed back into its home cage containing pre-weighed standard chow pellets, and the amount of chow consumed over the next 10 min was recorded.

### Tail suspension test

Approximately 4.5 weeks following the last alcohol exposure, mice underwent the tail suspension test (TST). Each mouse was hung by its tail for six min, and an experimenter blind to condition hand scored the last four min for active coping (hindlimb kicking, body shaking, attempts by the mouse to reach its tail or the wall) or passive coping (immobility and minor hindlimb movements) using BORIS (Behavioral Observation Research Interactive Software).

### Forced swim test

Approximately 5 weeks following last alcohol exposure, for the forced swim test (FST), each mouse was placed in a glass cylinder (1.5 L beaker, 15 cm h, 11 cm d) containing room temperature water for six min. The last four min were scored for swimming, climbing, and immobile behaviors by an experimenter blind to condition using BORIS. Swimming was scored as forceful movements of the hindlimbs. Climbing was scored as a vertical/tilted position against the cylinder wall surface with the head pointed upwards and vigorous kicking. Minor movements in the forelimbs and light kicking of one hindlimb to stay afloat were scored as immobility.

### Paw withdrawal

Approximately 5.5 weeks following the last alcohol exposure, mice were tested for pain sensitivity using a hot plate paw withdrawal assay. Mice received three trials separated by 10 min, in which they were placed within a cylinder on a hot plate of a temperature of 52 °C or 58 °C, and the latency to lift or lick its hind paw or jump away from the hotplate was recorded and the mouse was removed from the hot plate and returned to its home cage. If the mouse did not withdraw its paw within 30 sec, it was removed from the hot plate to prevent tissue damage. Paw withdrawal latency averaged across the three trials was calculated.

### Statistical analysis

Statistical analyses were performed in GraphPad Prism 9. Data for all dependent measures were examined for normal distributions within group and equality of variance across groups using Q-Q plots. For alcohol and water consumption, outliers for 2 hr, 4 hr, and 24 hr time points were determined using distributions of Day 4 total volume intake across all cycles for H20, 10%, and 20% EtOH DID groups within sex plotted together on Q-Q plots, and corresponding EtOH and H2O consumption values were excluded for analyses from the time point the outlier was identified through the rest of Day 4 time points (a total of 4 instances). For cumulative consumption only, excluded values were replaced with interpolated values based on the average of the mouse’s two adjacent data points (e.g., average of cycles 2 and 4 for interpolated cycle 3 data point). Analysis of variance (ANOVA) was used to evaluate the effects of and interactions between sex, EtOH concentration, and DID cycle on EtOH/water consumption and preference and between sex and EtOH on behavioral measures in protracted abstinence. For all ANOVAs, significant main effects and interactions were further probed with *post hoc* paired or unpaired t-tests with Holm-Sidak corrections for multiple comparisons to address a priori questions about sex differences in patterns of drinking and the effects of alcohol drinking and abstinence on behavior, and multiplicity-adjusted P values are reported. Statistical comparisons were always performed with an alpha level of 0.05 and using two-tailed analyses. Data are presented as mean ± SEM, and raw data points are included in all graphs where reasonable.

## Results

### Distinct patterns of alcohol and water consumption depending on sex and alcohol concentration

Here we used a modified DID binge drinking paradigm to assess the effects of sex on water and alcohol consumption and preference across multiple concentrations of alcohol (**Fig. 1a**). We first measured raw cumulative intake of EtOH solution or H2O in the 20% EtOH DID, 10% EtOH DID, and H2O DID mice (**Fig. 1c-e**). A 2xRM-ANOVA on raw cumulative intake of 20% EtOH showed main effects of sex (F (1, 18) = 5.97, *P = 0.025) and cycle (F (7, 126) = 1141.0, ****P < 0.0001) and an interaction between the two (F (7,126) = 4.45,***P = 0.0002). Post hoc t-tests with H-S corrections showed that raw intake volume was higher in F than M on cycles 5-8 (cycle 5: t144 = 2.67, *P = 0.050; cycle 6: t144 = 3.08, *P = 0.019; cycle 7: t144 = 3.60, **P = 0.005; cycle 8: t144 = 3.5, **P = 0.006; all other cycles: Ps > 0.30; **Fig. 1c**). In contrast, cumulative raw intake of 10% EtOH was not different in M and F, as there was an effect of DID cycle (F (7,126) = 702.30, ****P < 0.0001) but no effect of sex or interaction (Ps > 0.5) in a 2xRM-ANOVA (**Fig. 1d**). And, cumulative raw intake of water was not different between M and F in H2O DID (2xRM-ANOVA: main effect of DID cycle (F (7,126) = 311.90, ****P< 0.0001) but no effect of sex or interaction (Ps > 0.40; **Fig. 1e**). These results show that raw solution intake was greater in F than M only for high concentration (20%) EtOH. When EtOH or water consumption was normalized to bodyweight, females cumulatively consumed more 20% EtOH, 10% EtOH, and H2O than M (**Fig. 1f-h**). This emerged by cycle 3 for 20% EtOH DID, cycle 4 for 10% EtOH DID, and cycle 6 for H2O DID. A 2xRM-ANOVA for 20% EtOH DID showed main effects of sex (F (1, 18) = 29.37, ****P < 0.0001) and cycle (F (7,126) = 862.3, ****P < 0.0001) and an interaction between the two (F (7,126) = 28.42, ****P < 0.0001). Post hoc t-tests with H-S corrections showed that normalized consumption was higher in F than M on cycles 3-8 (*Ps < 0.05, as indicated; cycles 1 and 2: Ps > 0.15; **Fig. 1f**). A 2xRM-ANOVA for 10% EtOH DID showed main effects of sex (F (1, 18) = 14.11, **P = 0.0014) and cycle (F (7,126) = 810.6, ****P < 0.0001) and an interaction between the two (F (7,126) = 16.26, ****P < 0.0001). Post hoc t-tests with H-S corrections showed that normalized consumption was higher in F than M by cycles 4-8 (*Ps < 0.05, as indicated; cycles 1-3: Ps > 0.10; **Fig. 1g**). A 2xRM-ANOVA for H2O DID show main effects of sex (F (1, 18) = 9.73, **P = 0.006) and cycle (F (7,126) = 368.3, P < 0.0001) and an interaction between the two (F (7,126) = 10.42, ****P < 0.0001). Post hoc t-tests with H-S corrections showed that normalized H2O consumption was higher in F than M by cycles 6-8 (*Ps < 0.05, as indicated) and trended toward higher on cycle 5 (^$^P = 0.062; **Fig. 1h**), suggesting that females drank more (cumulative) water than males across the entire mDID paradigm, but increased bodyweight-normalized consumption was more robust for increasing EtOH concentrations. Examination of the average EtOH/water consumption for the first 2 hrs/day on each DID cycle (average of days 1-3 and the first 2 hrs on day 4) showed that females consumed more EtOH than males but not more water than males during this early access period (**Fig. 1i-k**). A 2xRM-ANOVA for 20% EtOH DID showed main effects of sex (F (1, 18) = 24.22,***P = 0.0001) and cycle (F (7,126) = 6.604, ****P < 0.0001) and an interaction between the two (F (7,126) = 2.12, *P = 0.046). Post hoc t-tests with H-S corrections showed that average 2 hr consumption was higher in F than M on cycles 2-8 (*Ps < 0.05, as indicated; cycle 1: P = 0.141; **Fig. 1i**). For 10% EtOH DID, a 2xRM-ANOVA showed main effects of sex (F (1, 18) = 18.19, ***P = 0.0005) and cycle (F (7,126) = 2.09, *P = 0.049) but no interaction between the two (P = 0.1707). Post hoc t-tests with H-S corrections showed that average 2 hr consumption was higher in F than M on cycles 3, 4, and 6 (*Ps < 0.05, as indicated) but not other cycles (Ps>0.14; **Fig. 1j**). For H2O DID, a 2xRM-ANOVA showed a main effect of cycle (F (7,126) = 4.66, ****P = 0.0001) but no effect of sex or interaction (Ps > 0.25; **Fig. 1k**). These results confirm that females’ increased consumption of EtOH compared to males was more robust for 20% than 10% EtOH. Further, they suggest while females display higher cumulative fluid consumption across the mDID paradigm for all EtOH/H2O conditions (**Fig. 1f-h**), their higher consumption of EtOH during the first two hr of each drinking day (**Fig. 1i,j**) was not attributable to higher fluid consumption, including water, during that time (**Fig. 1k**).

To further characterize drinking patterns, we next dissected Day 4 two bottle choice (2-BC) drinking behavior, examining both consumption and preference across the 2 hr, 4 hr, and 24 hr time points (**Figs. 2 and 3**).During the first 2 hrs of Day 4 of 20% EtOH DID, females drank more EtOH but not water than males (**Fig. 2a**). A 2xRM-ANOVA for EtOH consumption showed main effects of sex (F (1, 18) = 16.29, ***P = 0.0008) and cycle (F (7,126) = 5.72, ****P < 0.0001) but no interaction between the two (P = 0.268). Post hoc t-tests with H-S corrections showed that Day 4 2-hr consumption was higher in F than M on cycles 2, 5, and 7 (*Ps < 0.05, as indicated) and trended toward an increase on cycles 4, 6, and 8 (^$^Ps < 0.10) but did not differ on cycles 1 and 3 (Ps > 0.30; **Fig. 2a**). In contrast, water consumption did not differ between females and males, as there was a main effect of cycle (F (7,126) = 4.25, ***P = 0.0003) but no effect of sex or interaction (Ps > 0.25). Four-hr consumption showed a similar pattern (**Fig. 2b**), with a 2xRM-ANOVA for 4-hr 20% EtOH showing main effects of sex (F (1, 18) = 24.75, ****P < 0.0001) and cycle (F (7,126) = 4.13, ***P = 0.0004) but no interaction between the two (P = 0.714). Post hoc t-tests with H-S corrections showed that 4-hr consumption was higher in F than M on cycles 2, 3, 4, 5, and 8 (*Ps < 0.05, as indicated) and trended toward an increase on the remaining cycles (^$^Ps < 0.10). In contrast, for water consumption there was a main effect of cycle (F (7,126) = 4.03, ***P = 0.0005) and a trend for a main effect of sex (F (1, 18) = 3.74, ^$^P = 0.069) but no interaction (P > 0.55). For 24-hr 20% EtOH consumption (**Fig. 2c**), a 2xRM-mixed effects model showed main effects of sex (F (1, 18) = 28.12, ****P < 0.0001) and cycle (F (7, 116) = 2.40, *P = 0.025) but no interaction between the two (P = 0.495). Post hoc t-tests with H-S corrections showed that 24-hr consumption was higher in F than M on cycles 2-5 (*Ps < 0.05, as indicated) and trended toward an increase on cycle 7 (^$^P = 0.087). There was a trend towards a main effect of sex on water consumption (F (1, 18) = 4.11, ^$^P = 0.058) as well as a main effect of cycle (F (7, 116) = 5.06, ***P < 0.0001) but no interaction between the two (P > 0.50 at this timepoint. During the first 2 hrs of Day 4 of 10% EtOH DID, females drank more EtOH but not water than males (**Fig. 2d**). A 2xRM-ANOVA for EtOH consumption showed a main effect of sex (F (1, 18) = 21.77, ***P = 0.0002) and a trend toward a main effect of cycle (F (7, 124) = 2.0, P = 0.061) but no interaction between the two (P = 0.619). Post hoc t-tests with H-S corrections showed that Day 4 2-hr consumption was higher in F than M on cycles 2, 3, 4, and 7 (*Ps < 0.05, as indicated) but did not differ on cycles 1, 5, 6, and 8 (Ps > 0.15). In contrast, water consumption did not differ between females and males, as there was a trend toward an effect of cycle (F (7, 124) = 1.83, ^$^P = 0.0871) but no effect of sex or interaction (Ps > 0.40). Four-hr 10% EtOH consumption showed a similar pattern (**Fig. 2e**), with a 2xRM-ANOVA for 4-hr 10% EtOH showing a main effect of sex (F (1, 18) = 28.23, ****P < 0.0001) and a trend toward the effect of cycle (F (7, 124) = 2.06, ^$^P = 0.053) and an interaction between the two (F (7, 124) = 2.29, *P = 0.0312). Post hoc t-tests with H-S corrections showed that 4-hr 10% EtOH consumption was higher in F than M on cycles 2, 3, 4, and 7 (*Ps < 0.05, as indicated) but not the other cycles (Ps > 0.10). In contrast, water consumption did not differ between females and males, as there were no effects or interaction (Ps > 0.10). For 24-hr 10% EtOH consumption (**Fig. 2f**), a 2xRM-ANOVA showed main effects of sex (F (1, 18) = 15.81, ***P = 0.0009) and cycle (F (7, 123) = 5.00, ****P < 0.0001) but no interaction between the two (P = 0.188). Post hoc t-tests with H-S corrections showed that 24-hr consumption was higher in F than M on cycles 2, 3, and 7 (*Ps < 0.05, as indicated) and trended toward an increase on cycles 6 (^$^P = 0.074) and 8 (^$^P = 0.072) but was not different on cycles 1, 4, and 5 (Ps > 0.20). For water consumption, there was a trend towards a main effect of sex (F (1, 18) = 3.63, ^$^P = 0.073) and a main effect of cycle (F (7, 123) = 3.00, *P = 0.006) but no interaction (P = 0.705).

**Figure 2:**
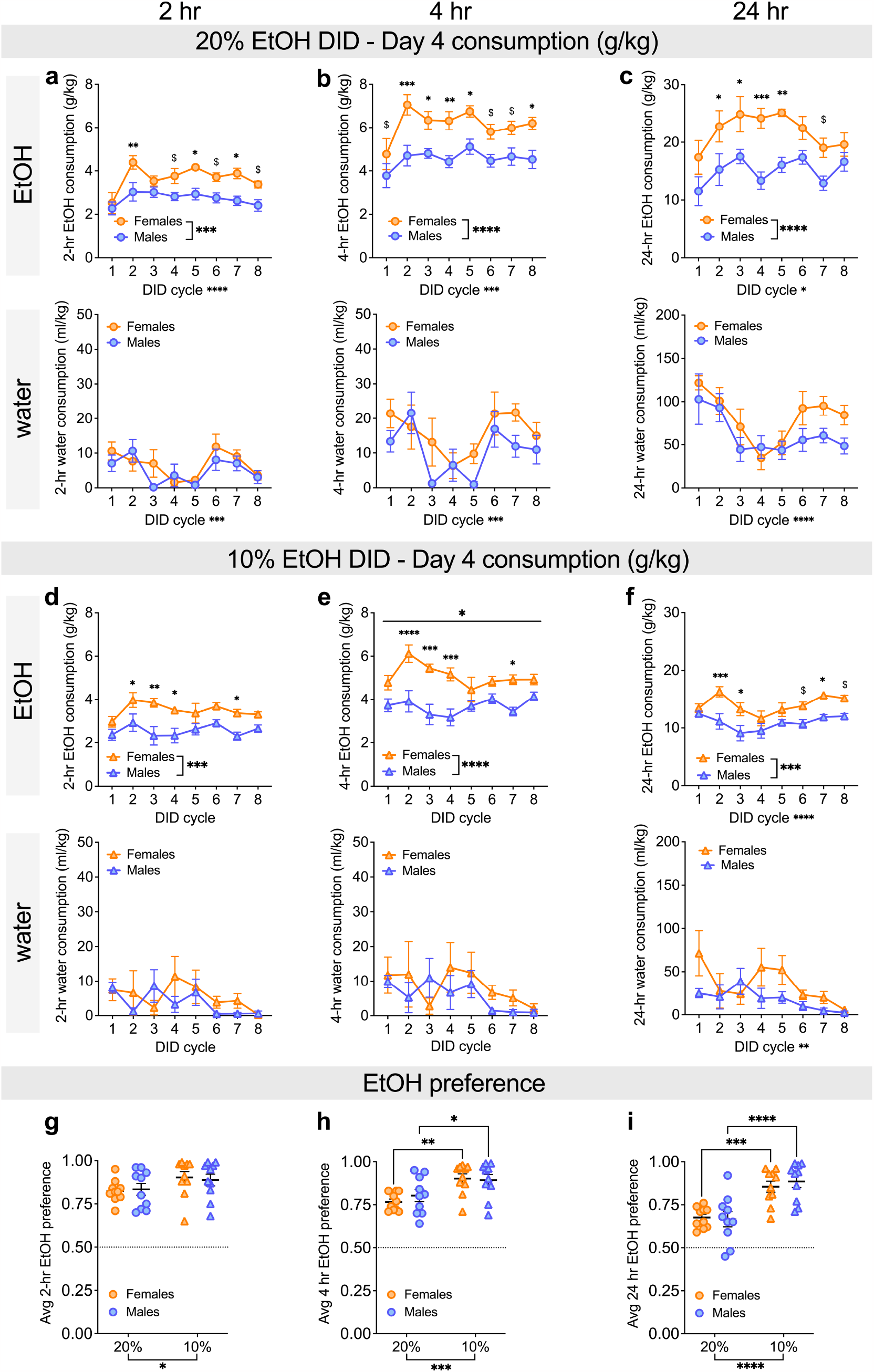
Females consume more alcohol than males across EtOH concentrations at 2-, 4-, and 24-hour timepoints on Day 4 access but have similar alcohol preference. **A-C)** EtOH (top row) and water (bottom row) consumption at 2-hr, 4-hr, and 24-hr time points on Day 4, 2-bottle choice (2-BC) access in 20% EtOH DID across cycles.**A)** During the first 2 hrs, females rapidly escalated EtOH drinking and consumed more 20% EtOH (g/kg) than males during DID cycles 2, 5, and 7, with a trend towards increased consumption on cycles 4 and 6. There was no difference in water consumption between the sexes. **B)** The sex difference in 20% EtOH consumption was even more robust for the first 4 hrs of Day 4 access, with females consuming more than males on DID cycles 2-5, and 8 and a trend towards increased consumption cycles 1, 6, and 7, with a trend towards a difference in water consumption between the sexes. **C)** Females consumed more 20% EtOH (g/kg) than males during the 24-hr period of Day 4 access on DID cycles 2-5, with a trend towards a difference in water between males and females. **D-F)** EtOH (top row) and water (bottom row) consumption at 2-hr, 4-hr, and 24-hr time points on Day 4, 2-bottle choice (2-BC) access in 10% EtOH DID across cycles.**D)** During the first 2 hrs, females consumed more 10% EtOH (g/kg) than males on DID cycles 3, 4, and 6, with a trend towards increased consumption in cycles 2 and 7. There was no difference in water consumption. **E)** During the first 4 hrs, females consumed more 10% EtOH (g/kg) than males during DID cycles 2, 3, 4, 6, and 7. There was no difference in water consumption. **F)** During the 24-hr access period, females consumed more 10% EtOH than males during DID cycles 2, 3, 6, and 7, with a trend toward increased consumption in cycle 8. Water consumption was overall higher in females compared to males. **G-I)** Average EtOH preference in 20% and 10% DID at 2-hr (**G**), 4-hr (**H**), and 24-hr (**I**) time points, showing that males and females similarly displayed higher EtOH preference for 10% compared to 20% EtOH. *P < 0.05,**P < 0.01, ***P < 0.001, ****P < 0.0001 in 2xRM-ANOVA main effects and interactions of sex and DID cycle (**A-F**) and 2xANOVA main effects of EtOH concentration (**G-I**) and post hoc t-tests with H-S corrections. ^$^P < 0.10 for post hoc t-tests with H-S corrections between M and F (**A-F**) and EtOH concentration within sex (**G-I**).

**Figure 3:**
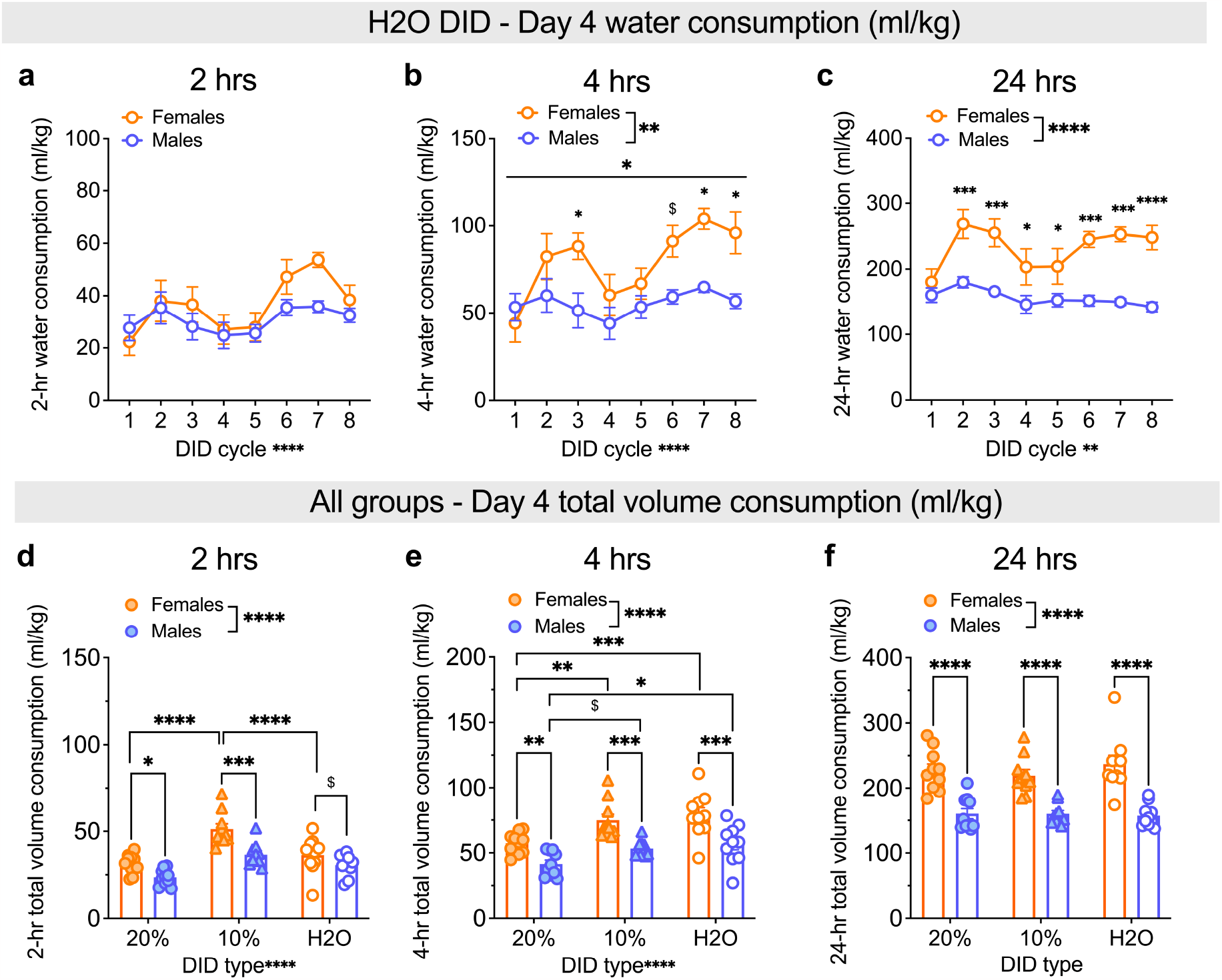
Higher female water consumption emerges across the day. **A-C)** Water consumption at 2-hr, 4-hr, and 24-hr time points on Day 4, 2-bottle choice (2-BC) access in H2O DID across cycles. **A)** There were no sex differences in H2O consumption (ml/kg) during the first 2 hrs of Day 4 access. **B)** Females consumed more H2O (ml/kg) than males during the first 4 hrs of Day 4 on cycles 3 and 8, with a trend towards increased consumption during cycle 6. **C)** Females consumed more H2O (ml/kg) than males during the 24-hr period on cycles 2-8. **D-F)** Average total volume consumption for Day 4 EtOH and water in 20% EtOH, 10% EtOH, and H2O DID. **D)** During the first 2 hrs, total volume consumption was highest for 10% EtOH, and females consumed more fluid than males in 10% EtOH DID. **E)** Within 4 hrs, females consume greater volume than males for all conditions, and consumption was higher for H2O, particularly in females. **F)** At the 24-hr time point, females consumed more fluid than males in all DID conditions; there were no differences in fluid consumption between DID conditions in either sex. *P < 0.05, **P < 0.01, ***P < 0.001, ****P < 0.0001 in 2xRM-ANOVA main effects and interactions of sex and DID cycle (**A-C**) and 2xANOVA main effects of sex and EtOH concentration (**D-F**), as well as post hoc t-tests with H-S corrections as indicated. ^$^P < 0.10 for post hoc t-tests with H-S corrections between M and F.

We evaluated whether sex or EtOH concentration affected EtOH preference at the 2, 4, and 24-hr time points on Day 4 2-BC (**Fig. 2g-i**). While EtOH preference was significantly above chance for both sexes at both 10% and 20% concentrations (one-sample t-tests compared to the null hypothesis of 0.5 preference score: all Ps < 0.01, not indicated), there were differences between concentrations that depended on sex. We found that 2-hr EtOH preference (averaged across all cycles) was higher for 10% than 20% EtOH (**Fig. 2g**). A 2xANOVA showed a main effect of EtOH concentration (F (1, 36) = 4.56, *P = 0.040) but no effect of sex or interaction (Ps > 0.65). However, post-hoc t-tests with H-S corrections showed no differences in preference for 10% versus 20% EtOH preference in either females (P = 0.156) or males (P = 0.2282). This phenotype was stronger at 4 hrs (**Fig. 2h**), as a 2xANOVA showed a main effect of EtOH concentration (F (1, 36) = 15.61, ***P= 0.0003) but no effect of sex or interaction (Ps > 0.40). Post-hoc t-tests with H-S corrections showed a higher 10% than 20% EtOH preference in females (t36 = 3.36, **P = 0.004) and males (t36 = 2.23, *P = 0.032). At 24 hrs, both sexes robustly displayed a stronger 10% than 20% EtOH preference (**Fig. 2i**). A 2xANOVA showed a main effect of EtOH concentration (F (1, 36) = 35.05, ****P < 0.0001) but no effect of sex or interaction (Ps > 0.55). Post-hoc t-tests with H-S corrections showed higher 10% than 20% EtOH preference in both sexes (F: t36 = 3.77, ***P = 0.0006; M: t36 = 4.61, ****P < 0.0001).

We also examined water consumption at these 2, 4, and 24 hr time points in mice that underwent H2O DID for comparison (**Fig. 3a-c**). For 2-hr consumption (**Fig. 3a**), a 2xANOVA revealed a main effect of cycle (F (7, 125) = 4.81, ****P < 0.0001) but no effect of sex or interaction (Ps > 0.15). For 4-hr H2O consumption (**Fig. 3b**), a 2xANOVA showed a main effect of sex (F (1, 18) = 11.86, **P = 0.003) and cycle (F (7, 125) = 4.92, ****P < 0.0001) and a trend for interaction between the two (F (7,126) = 2.20, *P = 0.039). Post hoc t-tests with H-S corrections showed that Day 4 4-hr consumption was higher in F than M on cycles 3, 7, and 8 (*Ps < 0.05, as indicated) and trended toward higher on cycle 6 (^$^P = 0.059) but did not differ on cycles 1, 2, 4, 5, or 7 (Ps > 0.25). For 24-hr H2O DID water consumption (**Fig. 3c**), a 2xRM-mixed effects model showed main effects of sex (F (1, 18) = 30.65, ****P < 0.0001) and cycle (F (7, 115) = 3.27, **P = 0.003) but no interaction between the two (P = 0.158). Post hoc t-tests with H-S corrections showed that 24-hr consumption was higher in females than males on all cycles but the first (cycle 1: P = 0.266; all other cycles: *Ps < 0.05, as indicated). These results show that water consumption in females is higher than in males, but this emerges across the 24-hr period after the first two hr of access. Given these results, we further assessed total volume consumed across 2, 4, and 24 hr time points of Day 4 averaged across cycles for all mice (**Fig. 3d-f**). For the 2-hr time point (**Fig. 3d**), a 2xANOVA showed main effects of sex (F (1, 54) = 24.46, ****P < 0.0001) and EtOH concentration (F (2, 54) = 25.23, ****P < 0.0001) but no interaction (P = 0.154). Post hoc t-tests with H-S corrections showed that total consumption for the first two hrs on Day 4 was higher in females than males for 10% (t54 = 4.42, ***P = 0.0001) and 20% EtOH (t54 = 2.39, *P = 0.04, with a trend towards a sex difference for H2O DID mice (P = ^$^0.085). Further, total volume consumed in 10% EtOH DID mice was higher than in 20% EtOH and H2O DID for females (***Ps ≤ 0.0001, as indicated) and than 20% DID mice for males (**P = 0.004; all other Ps > 0.10). For the 4-hr time point (**Fig. 3e**), a 2xANOVA showed main effects of sex (F (1, 54) = 39.94, ****P < 0.0001) and EtOH concentration (F (2, 54) = 12.71, ****P < 0.0001) but no interaction (P = 0.675). Post hoc t-tests with H-S corrections showed that total consumption for the first four hrs on Day 4 was higher in females than males for all three groups (**Ps < 0.01, as indicated). Further, total volume consumed in 20% EtOH DID mice was lower than in 10% EtOH and H2O DID for females (**Ps < 0.01, as indicated) and than in H2O DID mice for males (*P = 0.035), with a trend towards lower consumption than 10% EtOH (^$^P = 0.071). For the 24-hr time point (**Fig. 3f**), a 2xANOVA showed a main effect of sex (F (1, 54) = 87.29, ****P < 0.0001) but not EtOH concentration or interaction (Ps > 0.50). Post hoc t-tests with H-S corrections showed that total consumption for the first two hrs on Day 4 was higher in F than M for all three groups (****Ps < 0.0001). Thus, total volume intake converged across the last 20 hrs of access for all groups, leading to similar overall volume intake across DID conditions that was higher in females than males.

Because females’ greater alcohol consumption was apparent at 2 hrs while sex differences in water consumption did not emerge until later timepoints, we investigated the patterns of EtOH and H2O consumption (**Fig. 4**). We found that mice of both sexes drank more alcohol in the first two hours than the second two hours, but this was a more prominent effect in 10% compared to 20% EtOH. In 20% EtOH DID females (**Fig. 4a**), a 2xRM-ANOVA showed main effects of the first vs. second two hrs (“time”; F (1, 9) = 27.25,***P = 0.0005) and DID cycle (F (7, 63) = 2.97, **P = 0.009), as well as an interaction (F (7, 63) = 2.41, *P = 0.030). Post hoc t-tests with H-S corrections showed that consumption was higher on the first two compared to the second two hrs on cycles 2, 4, 5, 6, and 7 (**Ps < 0.01, as indicated) but not cycles 1, 3, or 8 (Ps > 0.15). We observed a similar, but more robust, pattern for 10% EtOH DID females (**Fig. 4b**). A 2xRM-ANOVA showed a main effect of time (F (1, 9) = 226.8, ****P < 0.0001) and DID cycle (F (7, 63) = 2.39, *P = 0.031) but no interaction (P = 0.149). Post hoc t-tests with H-S corrections showed that consumption was higher in the first two compared to the second two hrs on all cycles (***Ps < 0.001, as indicated). In contrast, water consumption for 20% and 10% EtOH females did not differ between the first vs. second 2 hrs of access on Day 4 or DID cycle, as 2xRM-ANOVAs showed no effects or interactions (all Ps > 0.10; **Fig. 4c**,**d**). In H2O DID females (**Fig. 4e**), a 2xRM-ANOVAs showed an effect of cycle (F (7, 63) = 5.33, ****P < 0.0001) and a trend for time (F (7, 63) = 5.33, ^$^P = 0.093) but no interaction (P > 0.20). These results suggest that females specifically frontload their consumption for EtOH, particularly at lower concentrations. While there was a trend for this front-loading behavior in water for females, it was less robust than alcohol consumption, likely because of the highly variable levels of water consumed. In concert with our results showing that total volume consumption is highest in 10% EtOH DID females in the first two hrs and then equilibrates by 24 hrs (**Fig. 3d-f**), alcohol frontloading may be performed to achieve similar levels to 20% EtOH consumption. Males also displayed frontloading behavior for EtOH, especially 10%, however, they also displayed a mild frontloading of H2O. In 20% EtOH DID males (**Fig. 4f**), a 2xRM-ANOVA showed main effects of time (F (1, 9) = 27.68, ***P = 0.0005) but not DID cycle or interaction (Ps>0.15) on EtOH consumption. Post hoc t-tests with H-S corrections showed that consumption was higher on the first two compared to the second two hrs on cycles 2, 3, 4, and 6 (*Ps < 0.05, as indicated) but not cycles 1, 5, 7, or 8 (Ps > 0.15). As in females, we observed a more robust pattern for 10% EtOH DID than 20% DID males (**Fig. 4g**). A 2xRM-ANOVA for EtOH consumption in 10% EtOH DID males showed a main effect of time (F (1, 9) = 79.30, ****P < 0.0001) but no effect of DID cycle or interaction (Ps > 0.20). Post hoc t-tests with H-S corrections showed that consumption was higher in the first two compared to the second two hrs on all cycles (**Ps < 0.01, as indicated). In contrast to females that only significantly frontloaded EtOH, males’ water consumption was modestly higher in the first two hrs compared to the second two hrs in 10% EtOH DID and H2O DID (but not 20% EtOH DID). A 2xRM-ANOVA for water consumption on 20% DID in males showed a main effect of cycle (F (7, 63) = 4.36, ***P = 0.0005) but no effect of time or interaction (Ps > 0.65; **Fig. 4h**). However, for 10% EtOH DID (**Fig. 4i**), there was a main of time (F (1, 9) = 5.13, *P = 0.0498) but not cycle or interaction (Ps > 0.05). Post hoc t-tests with H-S corrections showed that consumption trended toward being higher on the first two compared to the second two hrs on cycles 1 and 3 (Ps < 0.10) but no other cycles (Ps > 0.35). Similarly, for H2O DID (**Fig. 4j**), there was a main effect of time (F (1, 9) = 6.08, *P = 0.036) but not cycle or interaction (Ps > 0.30). Post hoc t-tests with H-S corrections showed no differences in the first two versus second two hrs at any DID cycle (Ps > 0.10).

**Figure 4:**
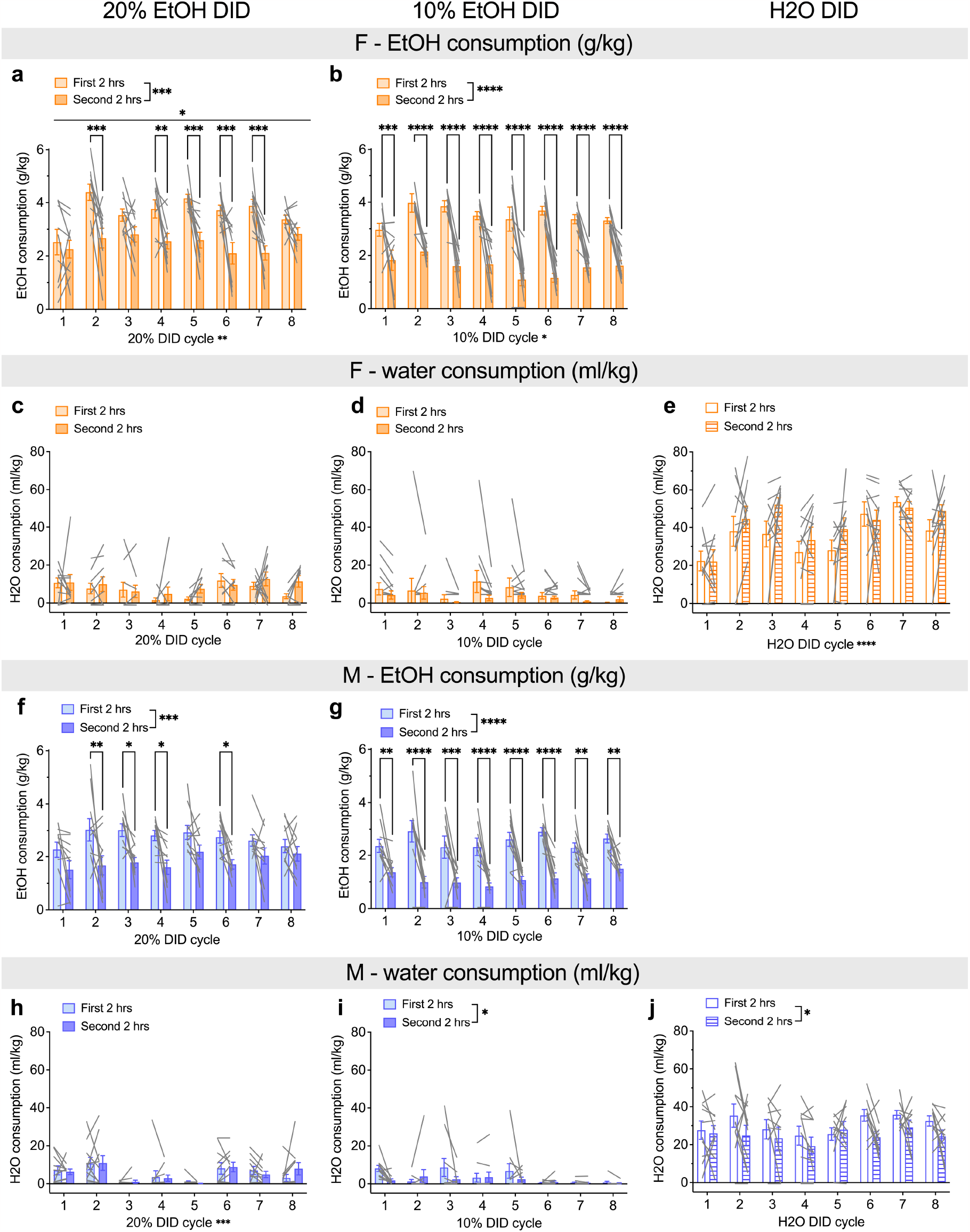
Both sexes display frontloading behavior in alcohol but not water consumption. **A-E**) EtOH and water consumption in females during the first vs. second 2 hrs of Day 4 for 20% EtOH, 10% EtOH, and H2O DID. **A)** Females consumed more 20% EtOH (g/kg) during the first vs. second 2 hrs of Day 4 DID on cycles 2 and 4-7. **B)** Females consumed more 10% EtOH (g/kg) during the first vs. second 2 hrs of Day 4 DID on all cycles. **C-E)** There was no difference in female H2O consumption (ml/kg) during the first vs. second 2 hrs in 20% EtOH DID (**C**), 10% EtOH DID (**D**), or H2O DID (**E**). **F-J)** EtOH and water consumption in males during the first vs. second 2 hrs of Day 4 for 20% EtOH, 10% EtOH, and H2O DID. **F)** Males consumed more 20% EtOH (g/kg) during the first vs. second 2 hrs of Day 4 DID on cycles 2-4 and 6. **G)** Males consumed more 10% EtOH (g/kg) during the first vs. second 2 hrs of Day 4 DID on all cycles. **H-J)** There was no difference in male H2O consumption (ml/kg) during the first vs. second 2 hrs in 20% EtOH DID (**H**), but consumption was higher on the first vs. second two hrs for 10% EtOH DID (**I**) and H2O DID (**J**). *P < 0.05, **P < 0.01, ***P < 0.001, ****P < 0.0001 in 2xRM-ANOVA main effects and interactions of time point and DID cycle, as well as post hoc t-tests with H-S corrections between the first and second 2 hrs.

To dissect the relationship between alcohol and water consumption in males and females, we performed correlational analyses at the 2, 4, and 24-hr timepoints (**Fig. 5**). We found that total volume intake was positively correlated with alcohol consumption in the 20% EtOH female and 10% EtOH male groups at the 2-hr (**Fig. 5a**; 20% EtOH F: R=0.89, ***P = 0.0006; 10% EtOH M: R=0.90, ***P = 0.0004; all other P > 0.14) and 24-hr (**Fig. 5c**; 20% EtOH F: R=0.69, *P = 0.0271; 10% EtOH M: R=0.81, **P = 0.0048; all other P > 0.15) timepoints. At the 4-hr (**Fig. 5b**) timepoints there was a positive correlation between alcohol consumption and total volume intake for all groups (20% EtOH F: R=0.79, **P = 0.0061; 20% EtOH M: R=0.65, *P = 0.0440; 10% EtOH F: R=0.66, *P = 0.0366; 10% EtOH M: R=0.76, *P = 0.0109). There was a trend towards a negative correlation between alcohol and water consumption in the 10% EtOH male cohort at 4 hrs (**Fig. 5b;** R=-0.63, ^$^P = 0.051) while there was a negative correlation at the 24-hr timepoint (**Fig. 5c**) in this group with no significant correlations in the other groups (R=-0.79, **P = 0.007; all other P > 0.15). Alcohol consumption and EtOH preference were positively correlated in the 4 (**Fig. 5b**; R=0.75, *P = 0.0132) and 24-hr (**Fig. 5c**; R=0.83, **P = 0.0027) 10% EtOH males only, with a trend at 2-hr (**Fig. 5a**; R=0.62, ^$^P = 0.056) for this group and in the 20% EtOH males at 24-hrs (R=0.62, ^$^P = 0.058; all other P > 0.18). These results suggest that males’ alcohol consumption was overall related to higher preference, while females’ alcohol consumption was unrelated to their water consumption at all time points.

**Figure 5:**
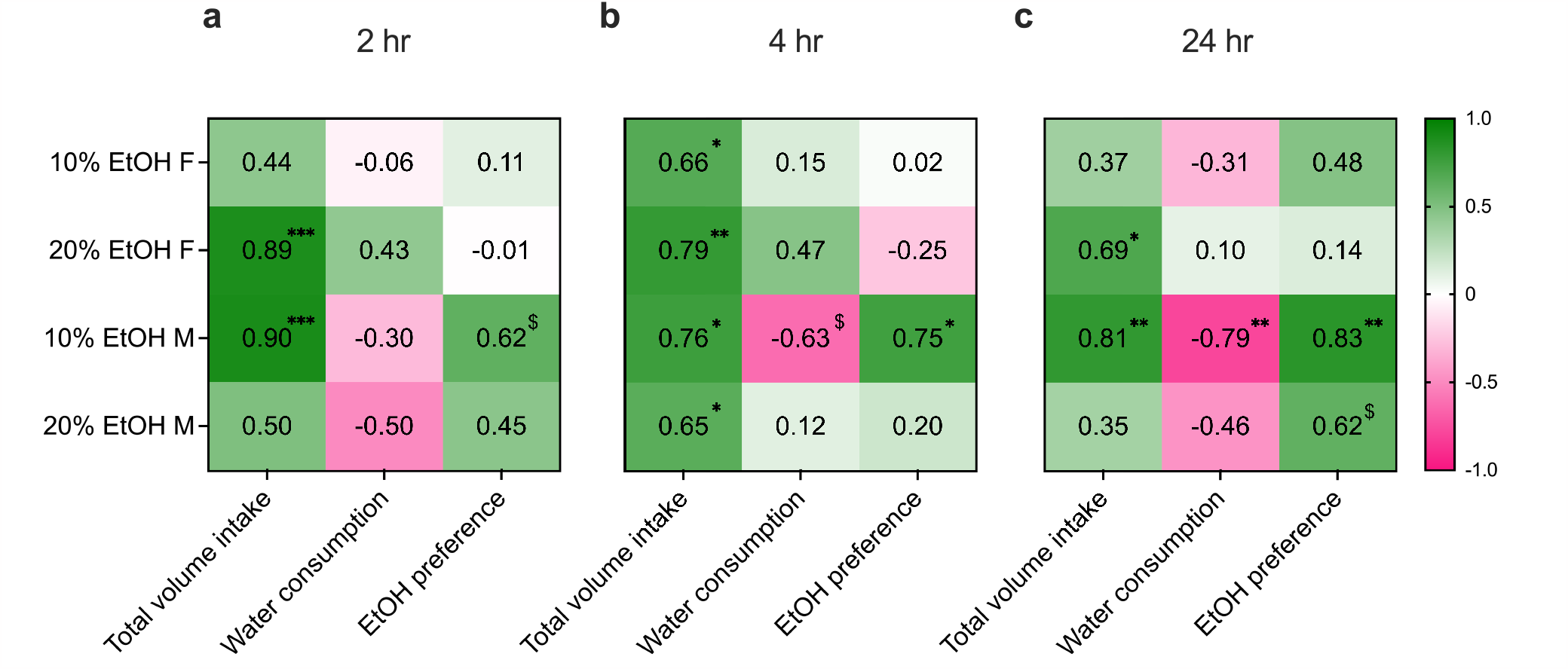
Alcohol consumption is correlated with water consumption and EtOH preference in males but not females. **A-C)** R values reported from linear regressions for correlations between alcohol consumption versus total volume intake, water consumption, and EtOH preference. **A)** At the 2-hr timepoint, there was a positive correlation between alcohol consumption and total volume in the 10% EtOH males and 20% EtOH females, and a trend towards a positive correlation between alcohol consumption and preference in the 10% EtOH males. **B**) At the 4-hr timepoint, alcohol consumption was positively correlated with total volume intake for both sexes in both the 20% and 10% EtOH cohorts, while there was a trend towards a negative correlation between alcohol and water consumption in the 10% EtOH male group and positive correlation between alcohol consumption and EtOH preference in this group only. **C**) At the 24-hr timepoint, alcohol consumption and total volume intake was positively correlated in the 20% EtOH female and 10% EtOH male groups. Alcohol and water consumption were negatively correlated while alcohol consumption and EtOH preference were positively correlated in the 10% EtOH male cohort only, with a trend towards a positive correlation between alcohol and EtOH preference in the 20% EtOH male group. *P < 0.05, **P < 0.01, ***P < 0.001 for correlations between alcohol consumption and total volume intake, EtOH preference, or water consumed at 2, 4 and 24-hr timepoints.

### No effect of chronic alcohol drinking on reward sensitivity

Following 20% EtOH or H2O DID, mice underwent a battery of behavioral assays to assess the effects of chronic binge alcohol drinking, beginning with sucrose preference starting 72 hrs after the last EtOH/water exposure (**Fig. 6**). Mice underwent 3 days of 1% sucrose preference, followed by 3 days of 2% sucrose preference, and we observed that females consumed more sucrose but not water than males at both doses, but EtOH drinking did not affect this behavior. A 3xRM-ANOVA on sucrose and water consumption for the 1% sucrose preference test revealed main effects of substance (F (1, 29) = 609.8, ****P < 0.0001) and sex (F (1, 29) = 48.38, ****P < 0.0001), as well as an interaction between the two (F (1, 29) = 20.03, ****P < 0.0001); however, there was no effect of or interactions involving EtOH as a variable (Ps > 0.20; **Fig. 6b**). To follow up on the substance x sex interaction, we performed 2xANOVAs within each substance. For sucrose, we found an effect of sex (F (1, 29) = 41.50, P < 0.0001) but no effect of EtOH or interaction between the two (Ps > 0.15). Post hoc t-tests with H-S corrections showed that CON and EtOH females consumed more sucrose than their male counterparts (t29 = 5.80, ****P < 0.0001; t29 = 3.43, **P = 0.002, respectively). We found similar results for 2% sucrose (**Fig. 6c**). A 3xRM-ANOVA showed main effects of substance (F (1, 35) = 931.1, P < 0.0001) and sex (F (1, 35) = 65.54, ****P < 0.0001), as well as an interaction between the two (F (1, 35) = 50.56, ****P< 0.0001); however, there was no effect of or interactions involving EtOH as a variable (Ps > 0.40). To follow up on the substance x sex interaction, 2xANOVAs within each substance were performed. For sucrose, we found an effect of sex (F (1, 35) = 60.76, P < 0.0001) but no effect of EtOH or interaction between the two (Ps > 0.55). Post hoc t-tests with H-S corrections showed that CON and EtOH F consumed more sucrose than their male counterparts (t35 = 5.86, ****P < 0.0001; t35 = 5.18, ****P < 0.0001, respectively). For water, we found no effects or interaction (Ps > 0.10). Analysis of preference for 1% sucrose over water showed no effects of sex or EtOH or an interaction between these variables (Ps > 0.25; **Fig. 6d**). Analysis of preference for 2% sucrose over water showed no effects of sex, EtOH, or an interaction between these variables (Ps > 0.10; **Fig. 6e**).Altogether, these results suggest that while females had higher consumption of sucrose compared to males, both sexes showed a high preference for sucrose at both concentrations, and these measures were not affected by a history of EtOH drinking.

**Figure 6:**
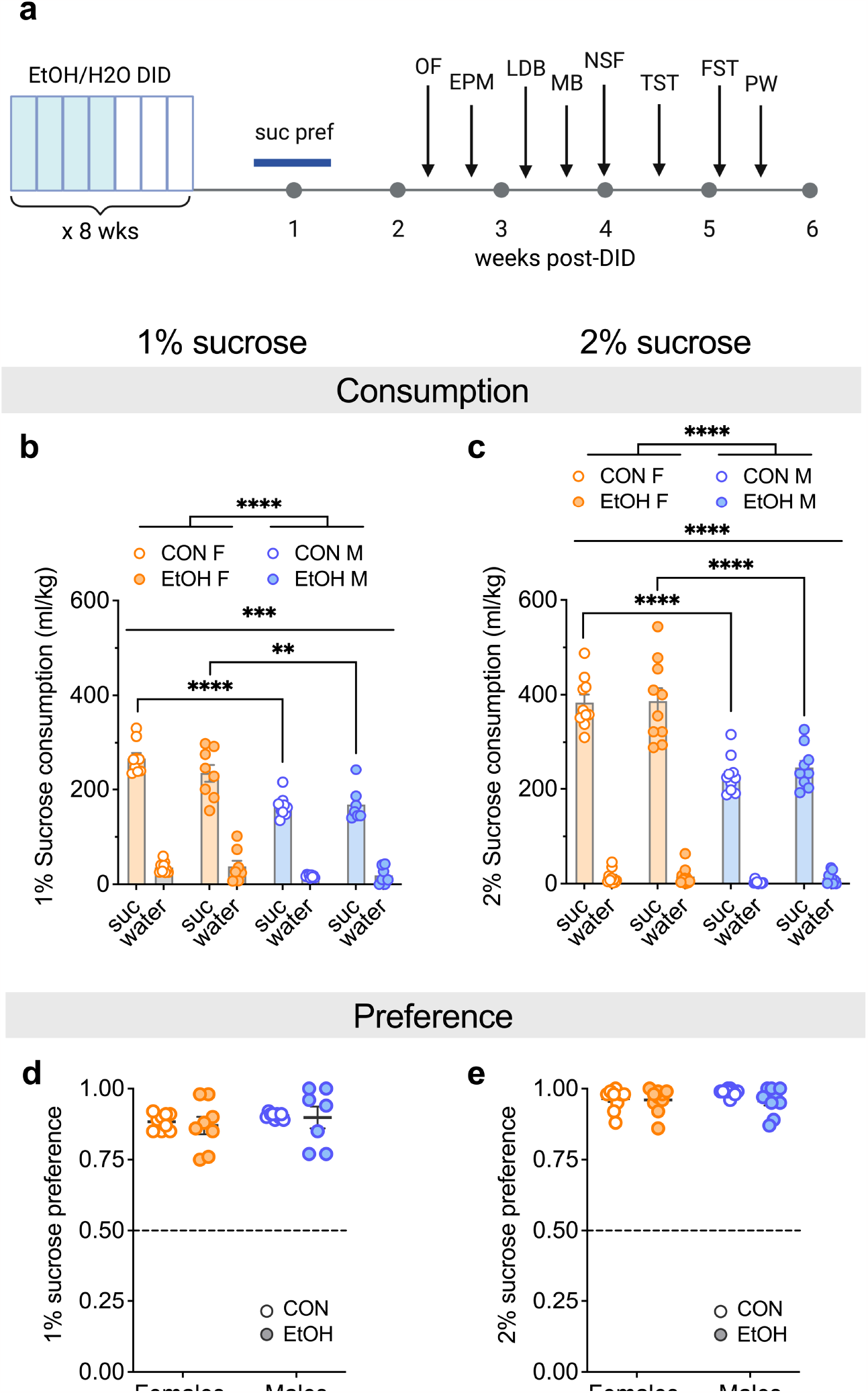
Females display higher sucrose consumption than males regardless of alcohol history. **A)** Experimental timeline. **B-C)** Sucrose and water consumption during the sucrose preference test, beginning 72 hr after the last EtOH/H2O drinking session, using 1% sucrose (**B**) and 2% sucrose (**C**), showing that females consumed more sucrose, but not water, than males, with no effect of EtOH drinking history. **D-E)** Sucrose preference was similar between H2O DID CON mice and 20% EtOH DID groups and between males and females for 1% (**D**) and 2% (**E**) sucrose. **P < 0.01, ***P < 0.001, ****P < 0.0001 in 3xRM-ANOVA main effects and interactions of substance, sex, and EtOH, as well as post hoc t-tests with H-S corrections as indicated.

### Behavioral disinhibition following a history of binge alcohol drinking was especially robust in females

We next assessed avoidance behavior across assays performed 2-4 weeks post-EtOH/H2O DID. We found that across behavioral tests, females displayed a reduction in avoidance behavior (**Fig. 7**). In the open field test (OF), 20% EtOH females, but not males, spent more time in the center of the OF than their H2O DID control (CON) counterparts (**Fig. 7a, b**). A 2xANOVA on the % time spent in the center showed a main effect of EtOH (F (1, 36) = 4.95, *P = 0.033) but no effect of sex or interaction (Ps > 0.15). Post hoc t-tests with H-S corrections showed that EtOH mice entered the center more quickly than CON mice in females but not males (F: t36 = 2.59, *P = 0.027; M: t36 = 0.56, ^$^P = 0.582). Examination of the % time in the center of the OF across the 30 min assay confirmed this effect (**Fig. 7b**). A 3xRM-ANOVA on the % time spent in the center of the open field revealed main effects of EtOH (F (1, 216) = 13.44, ***P = 0.0003) and time (F (5, 216) = 6.71, ****P < 0.0001), but no effect of sex (P > 0.40). However, there was a sex x EtOH interaction (F (1, 216) = 5.64, *P = 0.018) but no other interaction (Ps > 0.25). We followed up using 2xRM-ANOVAs within sex. In females, there were main effects of EtOH (F (1, 18) = 8.10, *P = 0.011) and time (F (5, 90) = 5.48, ***P = 0.0002), as well as an interaction between the two (F (5, 90) = 3.04, *P = 0.014). Post hoc t-tests with H-S corrections showed that EtOH females spent more time in the center in the 20-25 min time bin (t108 = 4.33, ***P = 0.002) and a trend in the 15-20 min time bin (t108 = 2.59, ^$^P = 0.053). In males, there was a main effect of time (F (5, 90) = 5.60,***P = 0.0002) but no effect of EtOH or interaction (Ps > 0.60). When the % time spent in the center of the OF was averaged across the 30 min test, a 2xANOVA showed a main effect of EtOH (F (1, 36) = 4.95, *P = 0.033) but no effect of sex or interaction (Ps > 0.15). Post hoc t-tests with H-S corrections showed that EtOH F spent more time in the center than CON F (t36 = 2.59, *P = 0.027) but there was no difference between CON and EtOH males (t36 = 0.56, P = 0.582). A 2xANOVA on the latency to enter the center of the OF showed a main effect of EtOH (F (1, 36) = 16.50, ***P = 0.0003) but no effect of sex or interaction (Ps > 0.20). Post hoc t-tests with H-S corrections showed that EtOH mice entered the center more quickly than CON mice in both sexes (F: t36 = 3.62, **P = 0.002; M: t36 = 2.12, *P = 0.041; **Fig. 7c**). A 2xANOVA on the total distance traveled during the assay showed no effects of EtOH or sex or interaction (Ps > 0.75; **Fig. 7d**), suggesting that differences in avoidance behavior were not due to locomotor effects.

**Figure 7:**
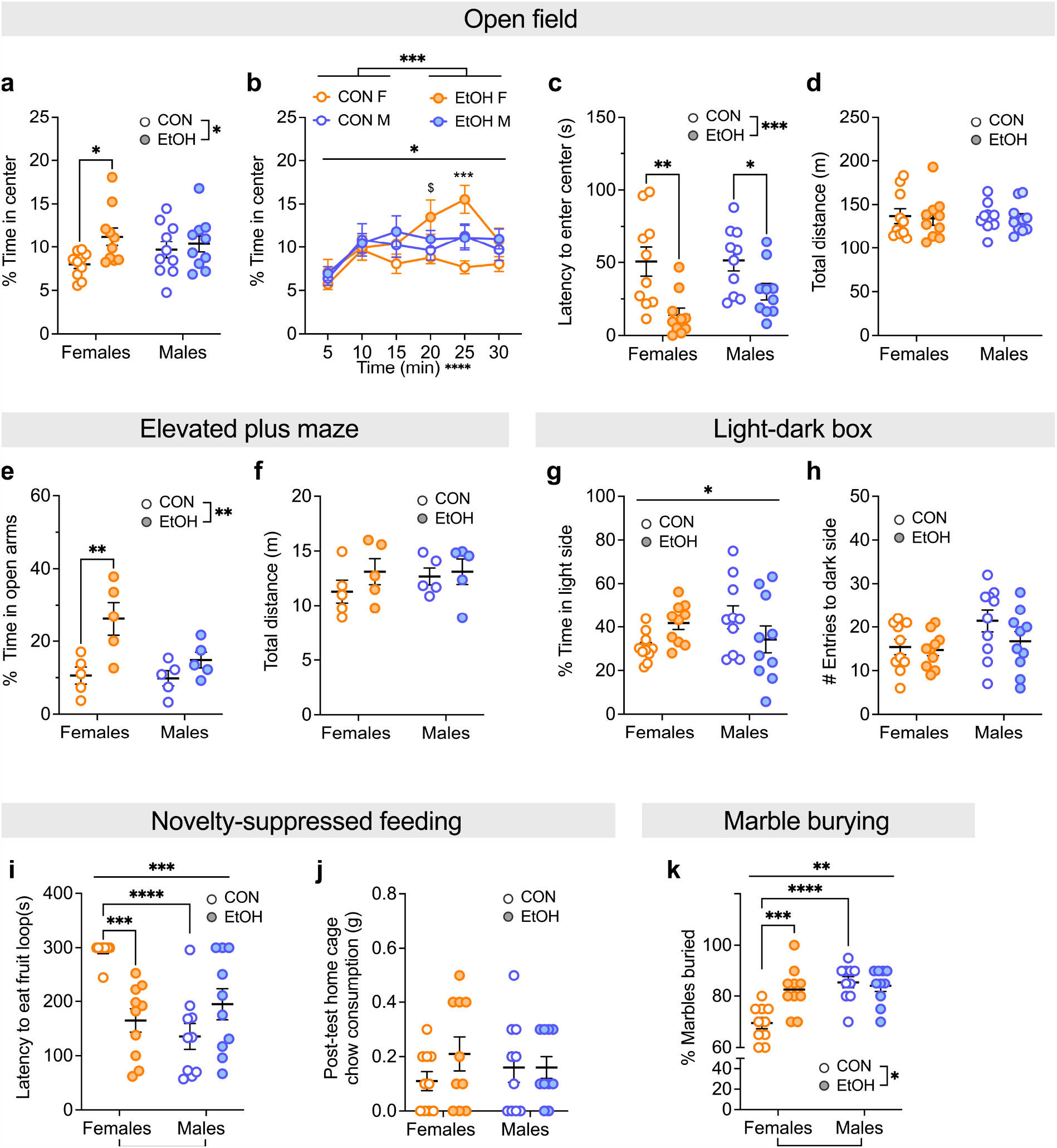
Females display reduced avoidance and increased compulsive behavior during protracted abstinence (2-4 weeks) from chronic binge alcohol drinking. **A-D)** Open field test (OF). **A)** EtOH females, but not males, spent a greater % time in the center of the OF compared to CONs. **B)** This effect emerged across the 30 min assay. **C)** EtOH mice of both sexes had a shorter latency to enter the center of the OF. **D)** There was no effect of sex or alcohol history on locomotion in the OF. **E-F)** Elevated plus maze (EPM). **E)** EtOH females, but not males, spent a greater % time in the open arms of the EPM compared to CONs. **F)** There was no effect of sex or alcohol history on total distance traveled (m) in the EPM. **G-H)** Light-dark box (LDB). **G)** There was an interaction between sex and alcohol history for % time spent in the light side of the LDB but no differences in direct comparisons. **H)** There was no effect of alcohol history or sex on locomotion as measured by the number of dark side entries. **I-J)** Novelty-suppressed feeding (NSF). **I)** EtOH females, but not males, had a reduced latency to eat the fruit loop compared to CONs. In addition, males had shorter latencies than females. **J)** There was no effect of sex or alcohol history on post-test home cage food consumption. **K)** Marble burying (MB). EtOH females, but not males, buried more marbles than CONs. In addition, males buried more marbles than females. *P < 0.05, **P < 0.01, ***P < 0.001, ****P < 0.0001 in 2xANOVA main effects of and interactions between sex and EtOH, 3xRM-ANOVA main effects of and interactions between sex, EtOH, and time, and post hoc t-tests with H-S corrections as indicated. ^$^P < 0.10 for post hoc t-tests with H-S corrections between M and F.

In the elevated plus maze (EPM), a 2xANOVA on the % time spent in the open arms showed a main effect of EtOH (F (1, 16) = 12.05, **P = 0.003), a trending effect of sex (F (1, 16) = 4.17, ^$^P = 0.058), and a trend for interaction (F (1, 16) = 3.21, ^$^P = 0.092). Post hoc t-tests with H-S corrections showed that EtOH females spent more time in the open arms of the EPM than CON females (t16 = 3.72, **P = 0.004) but there was no difference between CON and EtOH males (t16 = 1.19, P = 0.252; **Fig. 7e**). A 2xANOVA on the total distance traveled showed no effects of sex or EtOH or interaction between the two (Ps > 0.25; **Fig. 7f**), suggesting no differences in locomotion. In the light-dark box (LDB), a 2xANOVA on the % time spent in the light side showed no effect of sex or EtOH (Ps > 0.40) but there was an interaction between the two (F (1, 36) = 5.65, *P = 0.023). Post hoc t-tests with H-S corrections showed no effects in direct comparisons (Ps > 0.05; **Fig. 7g**). A 2xANOVA on the number of entries to the dark side, as a measure of locomotion, showed a trending effect of sex (F (1, 36) = 3.98, P = 0.054) but no effect of EtOH or interaction between the two (Ps > 0.15; **Fig. 7h**). In the novelty-suppressed feeding assay (NSF), a 2xANOVA on the latency to eat the fruit loop in the novel environment revealed an effect of sex (F (1, 36) = 8.90, **P = 0.005) but not EtOH (P > 0.10), and there was an interaction between the two (F (1, 36) = 19.06, ***P = 0.0001). Post hoc t-tests with H-S corrections showed that CON females had longer latencies than EtOH females (t36 = 4.24, ***P = 0.0003) and CON males (t36 = 5.20, ****P < 0.0001) but there was no difference between CON males and EtOH males (P > 0.30; **Fig. 7i**). A 2xANOVA on the post-test home cage chow consumption showed no effects of sex or EtOH or interaction between the two (Ps > 0.30; **Fig. 7j**), suggesting there were no differences in hunger driving effects of sex and EtOH on NSF. Altogether, results from avoidance behavior assays suggest that a history of binge alcohol drinking in females, but not males, leads to a decrease in behavioral inhibition. We also examined marble burying behavior, finding that EtOH females also display increased compulsive-like behavior (**Fig. 7k**), as a 2xANOVA on the proportion of marbles buried showed effects of sex (F (1, 36) = 13.46, ***P = 0.001) and EtOH (F (1, 36) = 5.81, P = 0.021) and an interaction between the two (F (1, 36) = 9.24, **P = 0.004). Post hoc t-tests with H-S corrections showed that CON females buried fewer marbles than EtOH females (t36 = 3.85, ***P = 0.001) and H2O M (t36 = 5.20, ****P < 0.0001) but there was no difference between CON and EtOH males (P > 0.30).

### Effects of sex, but not EtOH drinking, on coping strategy

Finally, we tested mice in the forced swim test (FST), tail suspension (TS), and paw withdrawal assay for pain 4-6 weeks following EtOH/H2O exposure (**Fig. 8**). We found that females displayed more active coping strategies, as a 2xANOVA on the % time immobile in the FST showed an effect of sex (F (1, 36) = 7.79, **P = 0.008) but no effect of EtOH or interaction between the two (Ps > 0.45). Post hoc t-tests with H-S corrections showed trends for greater immobility in males than females in CON and EtOH groups (CON: t36 = 1.93, ^$^P = 0.099; EtOH: t36 = 2.02, ^$^P = 0.099; **Fig. 8a**). A 2xANOVA on the % time swimming showed an effect of sex (F (1, 36) = 8.48, **P = 0.006) but no effect of EtOH or interaction between the two (Ps > 0.45). Post hoc t-tests with H-S corrections showed trends for greater swimming in female than male in CON and EtOH groups (CON: t36 = 1.95, P = 0.072; EtOH: t36 = 2.17, P = 0.072; **Fig. 8b**). A 2xANOVA on the % time climbing showed no effects of sex or EtOH or interaction between the two (Ps > 0.65; **Fig. 8c**). We also found more active coping in females than males on the TST, as a 2xANOVA on the % time passive showed an effect of sex (F (1, 36) = 9.09, **P = 0.005) but no effect of EtOH or interaction between the two (Ps > 0.15). Post hoc t-tests with H-S corrections showed that CON females displayed less time passive than CON males (t36 = 3.12, **P = 0.007) but there was no difference between EtOH F and EtOH M (P > 0.25; **Fig. 8d**). These results showed that females displayed more active coping while males showed more passive coping, but there were no effects of EtOH on this behavior. Finally, we measured thermal pain sensitivity, finding no effect of EtOH or sex at either low- or high-temperature stimuli (**Fig. 8e**). A 2xANOVA on paw withdrawal latency for 52 °C showed no effects of sex or EtOH or interaction between the two (Ps > 0.65). Similarly, at 58 °C there were no effects or interactions (Ps > 0.65), confirming that EtOH did not affect thermal pain sensitivity.

**Figure 8:**
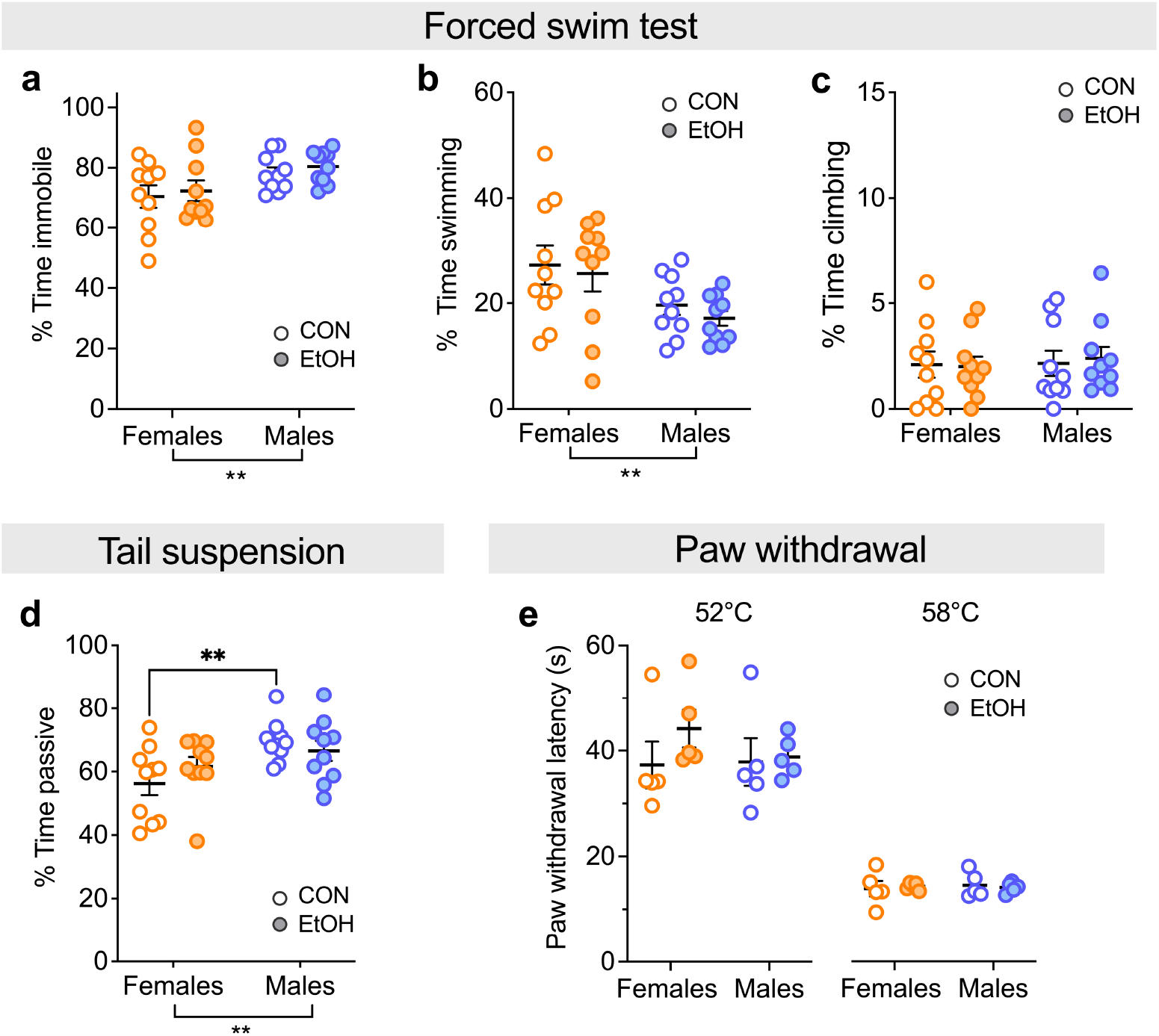
Sex, but not protracted abstinence from chronic binge drinking, impacts behavioral measures of coping strategy and negative affect tested 4-6 weeks following EtOH/H2O exposure. **A-C)** Forced swim test (FST). Females displayed less % time immobile (**A**) and more time swimming (**B**) but not climbing (**C**) than males, regardless of alcohol history. **D)** Tail suspension test (TST). Females displayed less % time passive than males. **E)** Hot plate paw withdrawal.There was no effect of sex or alcohol history on the paw withdrawal latency for either low (52 °C) or high (58 °C) hot plate temperatures. **P < 0.01 in 2XANOVA main effects of and interactions between sex and EtOH and post hoc t-tests with H-S corrections between M and F.

## Discussion

Here, we found that female mice consume more alcohol compared to their male counterparts in a modified DID paradigm across eight cycles (**Figs. 1 & 2**), consistent with what we and others have previously shown for binge alcohol drinking models in mice (8, 9). This difference was more robust for 20% compared to 10% EtOH, even though preference for alcohol over water did not differ between sexes at either alcohol concentration. Females consumed more alcohol than males across timepoints, an effect that was not dependent on overall higher fluid intake (**Fig. 5**). In addition, we found that protracted abstinence (2-4 weeks) from chronic binge drinking led to a behavioral disinhibition phenotype that was especially robust in females (**Fig. 7**). Specifically, we observed a reduction in anxiety-like and an increase in compulsive-like behavior in alcohol-abstinent females but not males compared to water control mice in the OF, NSF, and MB assays, as well as a stronger effect of alcohol experience for females in reducing avoidance behavior as measured through the EPM (**Fig. 7**). We additionally found basal sex differences in assays measuring reward sensitivity/anhedonia and coping behavior but no effects of alcohol abstinence in the sucrose preference, FST, and TS assays (**Figs. 6 & 8**). Together, these results suggest that anxiety and compulsive-related behaviors are most vulnerable to pathological behavioral outcomes in protracted alcohol abstinence from chronic binge drinking, especially in females.

We found that females consistently displayed increased alcohol consumption compared to their male counterparts across all timepoints (**Fig. 2A-F**). Females also consumed more water than males at most timepoints (**Fig. 3A-C**), an effect that is consistent with the literature and indeed, it has been previously reported that this may be dependent on estrogen fluctuation across the estrous cycle (29-32). While females with access to water only (H2O DID) consumed more water than males at the later timepoints (**Fig. 3B, C**), this effect became less robust when alcohol was available (**Fig. 2A-F**). Further, alcohol and water consumption in males was negatively correlated at the later timepoints (**Fig. 5B, C**) but for females there was no such relationship at any timepoint; others have similarly reported a dissociation between alcohol and water consumption in females (29). Altogether these data suggest that greater female alcohol consumption is not due to higher fluid consumption, and therefore that higher alcohol consumption in females is a factor of motivated drinking rather than thirst. EtOH preference, in contrast, was not different between the sexes (**Fig. 2-G-I**). At the 2-hr timepoint, this is likely due to the very low water consumption in both sexes that results in high preference for all mice; at 4 and 24 hrs, we conclude that the lack of sex difference in EtOH preference is due to the remaining trend in higher water consumption in females and that males’ alcohol consumption was correlated to their preference while females’ was not (**Fig. 5**). Overall, these results suggest a critical sex difference in the relationship between alcohol and water consumption: males’ alcohol consumption was overall positively related to their preference, and they tended to drink less water when they drank more alcohol (at least by 24 hrs). In contrast, females’ alcohol consumption was unrelated to their water consumption at all time points and therefore increased alcohol and water consumption are likely separate phenomena. One potential explanation for why female mice consume more alcohol than their male counterparts is that they have greater motivation for this behavior. Alternatively, females may require a higher dose to achieve BECs associated with the optimal interoceptive effects of alcohol. However, females across mammalian species, from mice to humans, display higher BECs for similar amounts of orally ingested alcohol. This is due to several body composition factors affecting alcohol pharmacokinetics, including lower water and higher fat content that affect volume distribution/absorption and lower gastric alcohol metabolism (30-33). Future studies will be able to directly determine the relationship between alcohol’s actions and metabolism in the periphery and the behavioral effects associated with alcohol drinking and abstinence. Overall, our findings highlight critical sex differences in motivated alcohol consumption and potential female vulnerability to the effects of high volume binge drinking, which is notable considering that binge drinking is particularly harmful to women across multifactorial negative health outcomes compared to men (7).

In this study, we found that the behavioral effects of protracted abstinence following chronic binge drinking were more robust in females, who demonstrate a behavioral disinhibition phenotype including reduced avoidance behavior and increased compulsive-like behavior (**Fig. 7**). These results underscore the importance of sex and timepoints into abstinence tested when investigating the effects of chronic alcohol on behavior.Indeed, the results of previous work on the affective behavioral consequences of chronic alcohol abstinence in C57BL/6J mouse models are variable as to time into abstinence tested and paradigms utilized, as well as highly male-dominated (16). Many studies have shown that males show increased avoidance, anhedonia, and compulsive behaviors in early abstinence (>1 week) following chronic alcohol across vapor inhalation (34-39), continuous access (21, 40), intermittent drinking (13), and DID (18, 22, 41, 42). Behavior during protracted abstinence (<1 week) in males is somewhat variable, with increased avoidance following chronic vapor (17) and DID (15, 18), with no effects following continuous (20, 21) and on most measures during intermittent access (10), and fairly consistent increased anhedonia across alcohol vapor (17, 19) and continuous drinking (20, 21, 40), with some exceptions for DID (18, 23). Notably, an increase in avoidance behavior during abstinence in males has been broadly observed in the literature (16), in contrast to our observation of a reduction in this behavior and one other study that showed mild behavioral disinhibition in males and females following seven weeks of intermittent alcohol access during early abstinence (24 hr since last alcohol consumption; (10). It is possible that the variability of avoidance-related alterations following chronic alcohol consumption in males is due to differences in testing timepoints.

Women show increased stress responsivity to various physiological triggers (43-45) and are more likely to use alcohol to regulate negative emotional states than men (46, 47), but there is a relative dearth of studies on the behavioral effects of chronic drinking in females rodent models. Most, however, point to no effect on avoidance in early or protracted abstinence on most measures (10, 34, 42), with some exceptions in continuous (48) and intermittent access (13) models. The literature on the effects of alcohol abstinence on anhedonia is limited, but investigations of protracted abstinence following chronic continuous access show an anhedonic phenotype in both sexes (20, 21, 40, 48-51). This is notable given our findings that protracted abstinence from binge drinking produces reduced avoidance with no changes to affect in females. Because these are some of the only studies investigating the long-term consequences of chronic alcohol exposure in females, future work directly comparing the nature of alcohol access, duration/amount of exposure, and specific behavioral time points will be useful for resolving which factors lead to specific behavioral outcomes in females. It is possible that this behavioral disinhibition phenotype is a protective effect in females, such that in our model female mice may be less prone to maladaptive anxiety-like behavior following chronic binge drinking. However, it is more likely that this reflects aberrant risk-taking behavior, given the established relationship between risk-taking/compulsivity, and alcohol drinking/relapse in humans and rodents (53-56).

Our results indicating sex differences in behavioral disinhibition during protracted abstinence led us to question whether the amount of cumulative alcohol consumption plays a role in the findings we observed here; that is, whether males would display a similar phenotype with additional alcohol drinking that achieved the cumulative alcohol consumption levels of females, or whether there are fundamental sex differences in the neural plasticity underlying these behavioral effects. Some studies report increased avoidance in males following acute abstinence from long term (8+ weeks) voluntary alcohol drinking (41) while others show no effect in some or all measures (42, 52) but to our knowledge, none have investigated these behaviors in protracted abstinence.

Future work is necessary to further our understanding of the mechanism underlying our finding that post-chronic alcohol behavioral disinhibition is more pronounced in females and to dissect the role of risk-taking and compulsivity in binge drinking behavior to refine our understanding of the results in our study. Thus, there are sex differences underlying the physiological and behavioral responses to alcohol, and women may be more vulnerable to maladaptive effects that potentially drive future risky behaviors.

### Perspective and Significance

Altogether, our results demonstrate sex differences in binge drinking and subsequent behavior in abstinence, pointing to the need for further investigations into the mechanisms underlying these phenomena.

Understanding anxiety-related behavior, anhedonia, and risk-taking during abstinence in both sexes is critical for informing knowledge on the effects of chronic binge drinking.

## Declarations

### Ethics approval and consent to participate

All animal procedures were performed in accordance with the Weill Cornell Medicine Institutional Animal Care and Use Committee guidelines and approval.

### Consent for publication

Not applicable.

### Availability of data and material

Source data will be publicly deposited upon publication.

### Authors’ contributions

K.E.P. designed all experiments, J.K.R.I., O.B.L., M.J.S, J.E.B, R.K., and K.E.P. performed and analyzed behavior data. K.E.P. oversaw experiments. K.E.P., L.J.Z., and T.B. wrote the manuscript, and all authors edited and approved the final version of the manuscript.

## Acknowledgements

Not applicable.

